# An integrated approach to comprehensively map the molecular context of proteins

**DOI:** 10.1101/264788

**Authors:** Xiaonan Liu, Kari Salokas, Fitsum Tamene, Markku Varjosalo

## Abstract

Protein-protein interactions underlie almost all cellular functions. The comprehensive mapping of these complex cellular networks of stable and transient associations has been made available by affinity purification mass spectrometry (AP-MS) and more recently by proximity based labelling methods such as BioID. Due the advancements in both methods and MS instrumentation, an in-depth analysis of the whole human proteome is at grasps. In order to facilitate this, we designed and optimized an integrated approach utilizing MAC-tag combining both AP-MS and BioID in a single construct. We systematically applied this approach to 18 subcellular localization markers and generated a molecular context database, which can be used to define molecular locations for any protein of interest. In addition, we show that by combining the AP-MS and BioID results we can also obtain interaction distances within a complex. Taken together, our combined strategy offers comprehensive approach for mapping physical and functional protein interactions.

## Introduction

Majority of proteins do not function in isolation and their interactions with other proteins define their cellular functions. Therefore, detailed understanding of protein-protein interactions (PPIs) is the key for deciphering regulation of cellular networks and pathways. During the last decade, the versatile combination of affinity purification and mass spectrometry (AP-MS) revolutionized the detailed characterization of protein complexes and protein-interaction networks^1^. The AP-MS approach relies on expression of a bait protein coupled with an epitope tag or antibodies targeting the endogenous bait protein, allowing purification of the bait protein together with the associating proteins (preys). This approach has been proven well suited for even large-scale high-throughput studies, and to yield highly reproducible data in both intra- and inter-laboratory usage^2^. The most commonly used epitope tags in medium to large-scale studies include FLAG^3^, His^4^, MYC^5^, HA^6^, GFP^7^and Strep^8^, of which the Strep-tag has become the gold-standard in affinity purification proteomics due to unparalleled protein purity in physiological purification conditions as well as the possibility for native competitive elution using biotin.

AP-MS can also be combined with quantitative proteomics approaches to better understand the protein complex stoichiometry^9^ and the dynamics of protein–complex (dis)assembly^1,10^. The combination of AP-MS with other techniques, such as biochemical fractionation, intact mass measurement and chemical crosslinking^11,12^, has been used to characterize supramolecular organization of protein complexes.

Although AP-MS remains the most used method for mapping protein-protein interactions, the recently developed proximity labeling approaches, such as BioID^13^ and APEX^14^, have become complementary and somewhat competing methods. BioID involves expression of the protein of interest fused with a prokaryotic biotin ligase (BirA) and the subsequent biotinylation of the amine groups of the neighboring proteins when excess of biotin is added to the cells. Whereas the wild-type BirA from *E. Coli* is capable of transferring the biotin only to a substrate bearing a specific recognition sequence, the generation of a promiscuous BirA* (Arg118Gly mutant) allows the biotinylation of any protein found within a 10 nm labeling radius^13,15^. While BioID has the abilities to capture weak and/or transient protein-protein interactions, the identified interactions are not limited to direct binders but can include proximate proteins as well.

In order to avoid artefactual interactions caused by overexpression of the bait proteins, majority of the large-scale interaction proteomic studies employ the Flp-In™ T-REx 293 cell line allowing moderate and inducible bait protein expression from isogenic cell clones^16^. Although the system allows rapid generation of transgene stably expressing cell lines, comprehensive analyses utilizing complementarily both AP-MS and BioID is resource-intense in the respect of cell line generation. To address this caveat and allow high-throughput comprehensive interactome analyses, we generated a *Gateway*^®^-compatible MAC (<u>M</u>ultiple <u>A</u>pproaches <u>C</u>ombined) -tag enabling both the single-step Strep AP-MS as well as the BioID analysis with a single construct, which decreases the number of required individual cell lines by 50%. In addition to allow visualization of tagged bait protein by immunohistochemistry, we included as well a nine amino acid hemagglutinin (HA)-epitope. The HA-epitope also facilitates additional follow-up approaches such as ChIP-Seq^17^ and purification of the crosslinked proteins for cross-linking coupled with mass spectrometry (XL-MS)^18^, making the MAC-tag almost as versatile as the Swiss Army knife.

To benchmark the usability and performance of the MAC-tag we applied it to 18 *bona fide* subcellular localization marker proteins. This allowed us to validate the correct localization of the MAC-tagged marker proteins as well as to monitor the localization of the *in vivo* biotinylated interactors. Additionally the interactions provide new information about these 18 marker proteins and their cellular functions. Furthermore the 18 localization markers and their 1911 interactions, form a basis of the reference molecular context repository, which we show can be used for “mass spectrometry (MS) microscopy” analysis of any protein of interest. The combinatory analysis with AP-MS and BioID also provided information, which efficiently could be used to derive relative spatial distances for proteins in a complex. Taken together, our devised combinatory MAC-tag and analysis approaches around it provide a plethora of information of the cellular functions and the molecular context of any studied protein.

## RESULTS

### MAC-tag AP-MS and BioID pipeline for detection of physical and functional interactions

To generate a versatile approach for identification of both stable physical and transient functional protein-protein interactions we integrated and optimized the BioID approach with our single-strep Strep AP-MS pipeline^10,23^. Both of these approaches have become the method of choice for interactomics analyses. We have recently shown the effectiveness of using these approached complimentarily^10,23^. However, the complementary use of the two techniques has been labor-intense, involving tagging of the bait proteins with BirA* and Strep-tag individually, as well as generation of two set of cell lines per bait. To overcome the major limitations, we have developed an integrated experimental workflow utilizing a MAC-tag containing both StrepIII-tag and BirA* (Supplementary Fig. 1a). In addition to optimizing the experimental steps, we focused on the compatibility of the two methods and to the simplicity of the analysis pipelines to generate a process with improved performance and reproducibility on detecting protein-protein interactions. The two pipelines differ only in the activation of the BirA* by addition of biotin to the cell culture media and harsher lysis condition in the BioID pipeline (Fig. 1, Supplementary Fig. 1). Without biotin addition the BirA* in the MAC-tag remains inactive (Supplementary Fig. 1b, c), resulting in identical (cor=0.88-0.99) single-step affinity purification results as vector with only StrepIII-tag (Supplementary Fig. 1d, e). Similarly, when biotin was added the results compare (cor=0.95-0.97) to that of a vector with BirA* alone (Supplementary Fig. 1d, e).

**Figure 1:**
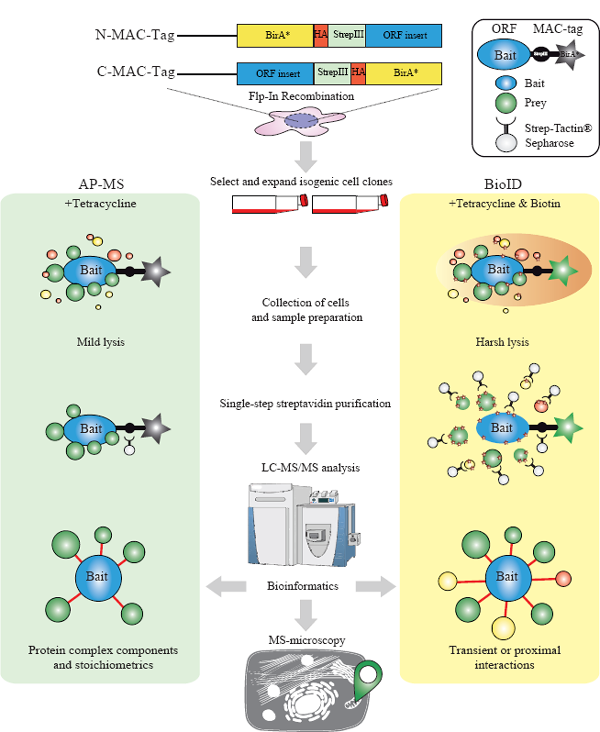
MAC-tag-based workflow for isolation and identification of protein complexes, protein-protein interactions and molecular context. Gateway compatible MAC-tag destination vectors containing Strepili, HA and BirA* were designed to allow the gene of interest either C- or N-terminal tagging. The expression vector can then be transfected into *Flp-In 293 T-REx* to establish the transgenic stably and inducible expressing isogenic cell lines. For the AP-MS and BioID analysis approaches, the cell line is separated into two cultures, BioID cells receiving addition of 50 μ M biotin in their culture medium. In the following protein extraction process, optimized lysis and affinity purification conditions for both analysis approaches were used. The interacting proteins were then analyzed by quantitative mass spectrometry and high confidence interaction proteins (HCIPs) were inferred via stringent statistical filtering. This integrated workflow allows laborless generation of cellular material for analyses, and results in integrated view of the formed protein complexes, protein-protein interactions and detailed molecular context definition.

The developed integrated approach significantly enhances (by two-fold) the throughput of generating bait-expressing cell lines, facilitates a comprehensive analysis of protein-protein interactions utilizing both the BioID and AP-MS, and allows analysis of protein complexes and even transient functional interaction networks with high sensitivity and reproducibility. Additionally the MAC-tag allows visualization of the bait protein with anti-HA antibody detecting the HA-epitope. This versatility of our approach was expected to give detailed view on the bait protein formed complexes, interactions, and actual molecular context via the detected stable, transient and/or proximal-interactions.

### Validation of the correct localization of the cellular localization markers and their in vivo biotinylated interactors.

We then went on and evaluated the MAC-tag system with 18 *bona fide* cellular localization markers (Supplementary Table 1) that cover most of cellular organelles to have more comprehensive view of the application of our integrated multiple approach system. Initially 18 localization markers were cloned to the MAC-tag vector, as a first step we explored their localization using fluorescence microscopy. The tagged-localization markers were visualized with anti-HA antibody and the *in vivo* biotinylated interactors with Alexa Fluor 594-Streptavidin (Fig. 2). These subcellular markers included: mitochondria (Apoptosis-inducing factor 1, AIFM1); endoplasmic reticulum (Calnexin, CALX); peroxisome (Catalase, CATA); early endosome (Early endosome antigen 1, EEA1); cytoplasmic peripheral plasma membrane marker (Ezrin, EZRI); nucleolus marker (rRNA 2′-O-methyltrans-ferase fibrillarin, FBRL); cis-Golgi marker (Golgin subfamily A member 2, GOGA2); chromatin (Histone H3.1, H31); exosome (Heat shock cognate 71 kDa protein, HSP7C); lysosome (Lysosome-associated membrane glycoprotein 1, LAMP1); nuclear envelope marker (Prelamin-A/C, LMNA); proteasome (Proteasome subunit alpha type-1, PSA1); recycling endosome (Ras-related protein Rab-11A, RAB11A); late endosome (Ras-related protein Rab-9A, RAB9A); microtubule (Tubulin alpha-1A chain, TBA1A) centrosome (Tubulin gamma-1 chain, TBG1); trans-Golgi (Trans-Golgi network integral membrane protein 2, TGON2); and ribosome (40S ribosomal protein S6, RS6) (Supplementary Table S1). All of the 18 MAC-tagged marker proteins localized to their corresponding cellular compartments, illustrating that the MAC-tag or the activation of the BirA* does not change the correct localization of these proteins. Furthermore, the localization of the *in vivo* biotinylated interactors correlates well with that of the corresponding localization marker. In addition to verifying the correct localization of marker proteins, the results highlight the usability of our MAC-tag constructs for fluorescence microscopy on detecting both the tagged-protein of interest as well as the interacting proteins.

### Identifying the physical and functional interactions of the cellular localization markers

Although many proteins and proteins families have been extensively studied with wide-range of cell biological or biochemical methods, others and we have shown the AP-MS and BioID can reveal wealth of new molecular and functional information^10,24^. However, for many proteins not much is known and there has not been systematic methods to efficiently and comprehensively characterize them. As shown in Fig. 2, the resolution of standard fluorescence microscopy does not allow capturing information of the protein dynamic localization and molecular context. Therefore, we MAC-tagged 18 known cellular localization markers and subjected them to our integrated method to obtain detailed molecular context proteome map with information from both the physical and functional interactions formed by these proteins. The analysis resulted in 26527 interactions from BioID and 9390 from AP-MS, of which 2118 high-confidence interactions from BioID and 679 interactions from AP-MS were retained after using stringent statistical filtering (Fig. 3a-c, Supplementary Table 1). The identified average connectivity (38) of the 18 localization markers, identified using AP-MS, matches well with the published large-scale studies^10,25^. As the BioID is also able to capture highly transient and close-proximity interactions, the total number of identified interactors as well as interactions per bait is higher than that of AP-MS (Fig. 3a, b). This is seen for example with Rab9A and Rab11A, two regulators of endosomal transport, for which BioID provides 16 times and 11 times more high-confidence interacting proteins (HCIPs) than AP-MS, respectively (Supplementary Fig. 2). In this case, the proteins detected solely with BioID likely represent cargo proteins in endosomal transit. Interestingly the ratio of identified novel vs. known interactions in total is almost two-fold higher with BioID (11.3) than with AP-MS (6.8), reflecting the sensitivity of BioID to identify more transient interactions (Fig. 3a).

**Figure 2:**
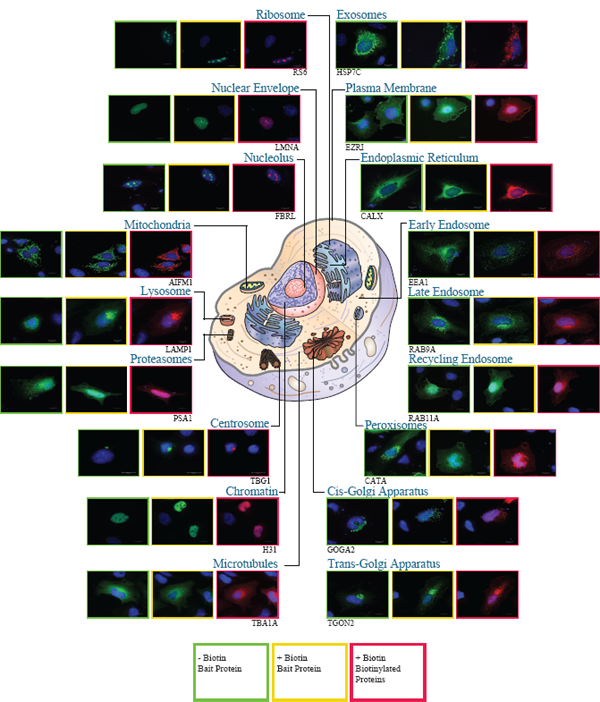
Fluorescence microscopy analysis of the bona fide cellular localization markers and their interactors. The 18 subcellular localization markers fused with MAC-tag were visualized by immunofluorescence staining using Alexa Fluor-488 labeled anti-HA immunostaining (green), their *in vivo* biotinylated interactors with Alexa Fluor-594 streptavidin (red), and cell nuclei with DAPI (blue) (Scale bar was 10**μ**m).

However, the complementary nature of these two methods is illustrated by their overlap as well as with their individually detected interactions, such as the ones formed by proteosomal marker PSA1^26^ and nuclear envelope marker LMNA (Fig. 3c, d). With PSA1 the overlap of AP-MS (green edges) and BioID (yellow edges) identified interactions is 17 components of the 20S core proteasome complex involved in the proteolytic degradation of most intracellular proteins (Fig. 3c, Supplementary Table 1c). BioID also captures myosins (MYH10 and MYH14) and unconventional myosins (MYO1B-D and −6), which have high turnover rates and after use they are either refolded for reuse or degraded by the proteasome ^27^. Additionally, BioID identifies proteasome activator complex subunits 1 (PSME1) and 2 (PSME2), which are part of the 11S (PA28) immunoproteasome^28^.

LMNA is a component of the nuclear lamina, playing an important role in nuclear assembly, chromatin organization, and framework for the nuclear envelope and telomere dynamics. Not surprisingly both the AP-MS and BioID identify interactions with lamin-B (LMNB)1, LMNB2, lamin-B receptor (LBR), lamina-associated polypeptide (LAP)2A and LAP2B, inner nuclear membrane protein Man1 (MAN1), and emerin (EMD). Another group of interacting proteins are nuclear pore complexes (NPCs) components: nuclear envelope pore membrane protein POM (P121A, P121C), Nuclear pore complex protein Nup (NU) 107, NU133, NU153, NU155, NU160, NUP50, NUP85, NUP98, NUP37, NUP43, nucleoporin SEH1, nucleoprotein TPR, ELYS, SEC13, of which only components of the nuclear basket NUP50, TPR, SEC13 are detected of low abundance with AP-MS (Fig. 3d, and Supplementary Table 1c). This suggests that for the correct localization to the nucleoplasmic side of the nuclear envelope, LMNA needs to pass through nuclear pore and during this process it transiently interacts and *in vivo* biotinylates the nuclear pore complex (NPCs) components. Similarly, the importin transport proteins importin subunit alpha (IMA1, IMA3, IMA4, IMA5, IMA6, IMA7), importin subunit beta-1(IMB1) are detected with BioID, but only IMA5 and IMA7 in AP-MS. In addition several histone modifiers and chromatin remodelers are detected (AN32E, EDF1, MYSM1 and PARP1). Somewhat surprisingly our analysis also identifies several proteins involved in cell cycle and mitosis (RAD50, ARF, KI67, HDGR2, WRIP1, AKP8L, BCCIP and P53).

Both of the examples show that the detected high-confidence interacting proteins are highly specific for the studied location, as illustrated by the retrieval of the HCIPs localization information from CellWhere database ^29^ (Fig. 3c, d). The proteins with the highest ranking for the particular location from CellWhere are shown in dark green and for the rest of the ranks the node color is light green. Proteins with no CellWhere ranking are shown in grey.

**Figure 3:**
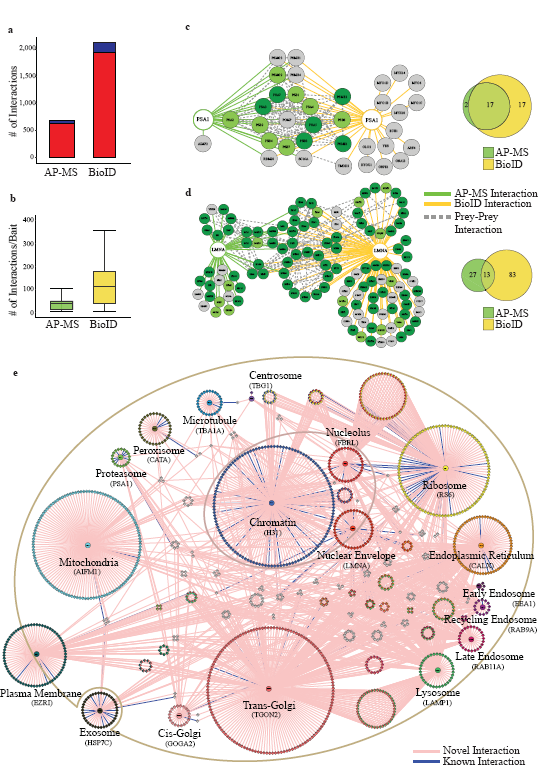
Generation of physical and functional PPIs interaction networks of bona fide cellular markers. The 18 localization markers we subjected to our integrated analysis, resulting in identification of 679 interactions from the AP-MS and 2118 interactions from BioID analysis. (**a**) The distribution of the number of known (blue) and novel (red) interactions within 18 *bona fide* subcellular organelle/structure markers, illustrate the need for systematic analyses. (**b**) The distribution of the number of interactions per localization marker by AP-MS or BioID purification approach shows similar distribution of connectivity as other publications using these approaches individually. (**c**, **d**) The protein-protein interaction network and molecular context for proteasome organelle marker (PSA1) and nuclear envelope (LMNA). The HCIs that were identified from AP-MS (green line) and BioID (yellow line) are shown together with the known prey-prey interactions (dashed grey line). The nodes are color-coded based on the localization rank obtained from the CellWhere database (key: dark green = primary cellular localization for the corresponding protein, light green = possible localization, grey = different or localization assigned for the protein). The Venn diagram highlights the complementary nature of the AP-MS and BioID approaches. (**e**) The reference molecular context map for the 18 subcellular organelles/structures. The unique high-confidence interactors from the BioID analysis are arranged in a circle around the corresponding localization marker and the shared interactors are shown with corresponding colors representing multiple localizations. Preys with more than four subcellular localizations are shown in white color. The novel interactions are shown in pink edges and the known interactions with blue.

### Reference molecular context proteome map reveals unique profiles for different cellular organelles

In addition to lacking molecular level resolution, standard fluorescence microscopy is often used to produce static images representing the particular time point when the image is taken. However, cellular proteins are highly diverse in their spatiotemporal properties, thus making their characterization with microscopy alone extremely challenging. The BioID, in principle, overcomes these limitations as monitoring of the biotinylated close-proximity proteins and their quantities should allow defining the BirA*-tagged bait proteins detailed molecular context within certain time period (Supplementary Fig. 2). Using the 2118 high-confidence interactions from BioID, we generated a cellular compartment-specific protein interaction map to the 18 *bona fide* localization markers (Fig. 3e and Supplementary Table1). The HCIPs domain as well as the gene ontology (GO) term profiles for each marker were unique (Supplementary Table 2 a-f and Supplementary Fig. 3, 4). However, we identified also shared HCIs between the endomembrane system consisting of ER (CALX), the Golgi (GOGA2 and TGON2), endosomes (EEA1, RAB9A and RAB11A) and lysosome (LAMP1). The four organelles shared 17 interactors, and the combination of any three locations shared in total 87 interactions (Supplementary Fig. 5a). These four organelles are involving in two major intracellular trafficking pathways: The exocytic pathway (ER via Golgi (53 shared interactions) to the plasma membrane); and the endocytic pathway (plasma membrane via endosomes to 1) Golgi (101 interactions) and 2) lysosome (61 interaction) to ER (86 interactions)). This organization is also well visible with, within a cell, the physically farthest from each other locating endosome and ER, sharing the least interactions of the all possible binary combinations of the four locations.

Similarly, chromatin, nucleolus and nuclear envelope are all sub-structures in the nucleus and shared interactors with each other (Supplementary Fig. 5b); nuclear envelope (LMNA) 22 with chromatin (H31) and chromatin 31 with nucleolus (FBRL). We previously already discussed the role of nuclear envelope on chromatin organization, and chromatin control the structure of nucleolus via ribosome DNA. Nucleolus is the place where ribosomal RNA transcription and the ribosome assembly occur. Ribosome (RS6) was detected as an outlier as it shared many of the interactors with other localization markers. This is explained by the fact that protein translation requires ribosomes, which are after the synthesis of the MAC-tagged protein immediately *in vivo* biotinylated in the BioID approach. Therefore the ribosome (RS6) was excluded from the further analyses. However, the other localization markers, such as mitochondria (AIFM1), cytoplasmic peripheral plasma membrane (EZRI), exosome (HSP7C), peroxisome (CATA), microtubule (TBA1A), proteasome (PSA1) and centrosome (TBG1) had highly unique molecular context signature, which suggest the usability of this reference set in tracking of protein of interests dynamic localization in intracellular environment. Additionally comparison of the HCIPs cellular locations from CellWhere database showed them to be assigned to the correct cellular localization according to their bait protein, further reinforcing the idea that proteins that share their interaction profiles are proximal. Finally, total of 1911 HCIs (excluding RS6 interactions), collapsed to 14 subcellular localizations were integrated to build up the reference molecular context map (Fig. 3e). Overlaying of any protein of interests BioID PPIs with our molecular map should allow defining the dynamic localization of the protein. In principle the developed MS-microscopy approach could have high impact on cellular quantitative biology (Fig. 4a).

**Figure 4:**
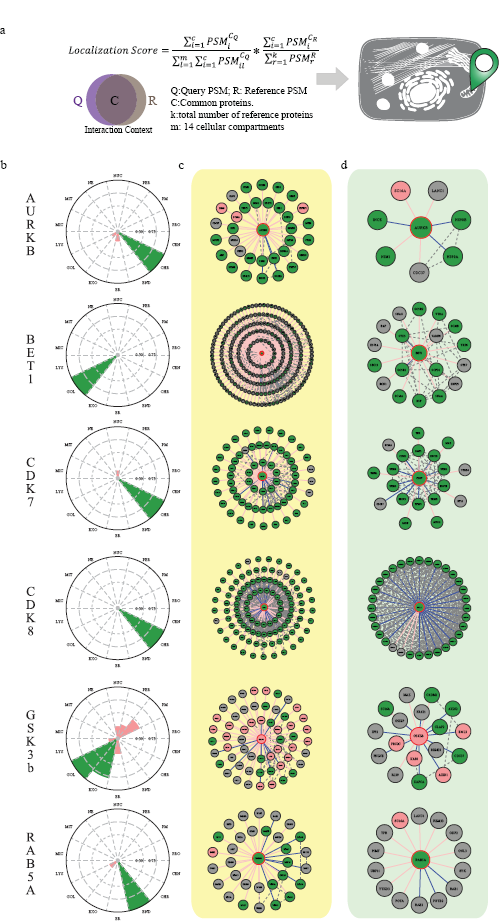
The reference molecular context map allows molecular level identification of cellular localization profiles for a protein of interest. (a) The schematic overview of the MS-microscope to assess the queried protein localization using our reference interaction context. (b) The polar plot shows the location of query protein observed by MS-microscopy. Each sector represents one subcellular location defined by our reference database. The color assigned to each of the localization is based on the annotation frequency (Pink: 0-0.5; Yellow: 0.5-0.75; Green: 0.75-1). (c,d) The PPIs interaction network obtained from BioID and AP-MS are shown separately. The localization of prey proteins was verified by Cellwhere database. Node color scheme coordinates the observation localization from Figure 4 b.

### Mass spectrometry microscopy using the reference molecular context proteome map

Despite the biological significance of dynamic subcellular localizations of proteins, simple tools for detecting the relative subcellular distribution have not been extensively developed. To test the applicability of the MS-microscopy on this, we selected dynamic cytoplasmic signaling molecules aurora kinase B (AURKB), cyclin-dependent kinase (CDK) 7, CDK8, and glycogen synthase kinase-*β* beta (GSK3B), as well as additional markers for cellular locations ras-related protein Rab-5A (RAB5A) and Golgi vesicular membrane-trafficking protein p18 (BET1) and applied our approach to them (Fig. 4, Supplementary Table 1a-c, and Supplementary Fig. 6). Aurora kinase plays an important role in cellular division by controlling chromatic segregation, which matches well to its interactions overlaying with “chromatin” marker H31 (Fig. 4b, c, d). Similarly SNARE protein BET1,involved in the docking process of ER-derived vesicles with the cis-Golgi membrane is assigned to Golgi (Fig. 4b, c, d). Essential component of the transcription factor II H (TFIIH), CDK7, and mediator complex associating CDK8 are predominantly associating with “chromatin” (Fig. 4b, c, d). This finding is in line with their important role in transcription regulation. Importantly, these examples show the high resolution of the MS-microscopy to distinct exact molecular locations, which could not be resolved by fluorescence microscopy (Supplementary Fig. 6). Glycogen synthase kinase β (GSK3B) phosphorylates many substrates in mammalian cells, and functions in many physiological processes, and acts as an important regulator in Wnt and Hedgehog signaling pathways^30^. Somewhat, to our surprise our MS-microscopy showed GSK3B localization to Golgi and exosomes. Recent research have demonstrated that a portion of GSK3B is localized to the trans-Golgi network through peripheral protein p230^31^ and that cytoplasmic GSK3B relocalizes to the same endosome as the internalized Wnt ligand^32^. It is plausible that this colocalization of GSK3B continues with active Wnt through endosomal organelles onto exosomes^33^. For validation of our endosomal location, we choose RAB5A which is known to localize to early endosomes and is involved in the recruitment of RAB7A and the maturation of these organelles to late endosomes^34^. In our analysis we can confirm the (early)-endosomal location as well as detect a fraction of Golgi localization, which could be related to the fusion of trans-Golgi network-derived vesicles with the early endosome^35^. These examples clearly establish the sensitivity and applicability of the MS-microscopy in defining the molecular context of a(ny) protein. In addition to analysis of wild type proteins our system should be useful for defining possible altered molecular context in human diseases caused by either somatic or germline genetic alteration, as well as for example analyzing functions of transgenic proteins not expressed in human cells.

### Transgenic Alternative Oxidase localizes in the mitochondria intermembrane and functionally associates with Complex I-V

Alternative oxidase (AOX) present in many lower eukaryotes, but not in vertebrates, transfer electrons directly from ubiquinol to oxygen in a nonproton-motive manner^36^. Transgenic expression of AOX in mammalian systems has been suggested as a therapeutic option for treating mitochondrial disease induced by OXPHOS dysfunction^37^, and additionally it has been shown that even broad expression of AOX does not disturb normal physiology in mice^38^. The exact molecular context of AOX in mitochondria membrane is not currently known, but it is thought to locate close to complex II based on its alleviating effects after toxic or pathological inhibition of the mitochondrial respiratory chain^39^. Therefore, we decided to apply our MS-microscopy method to define the molecular context of *Ciona intestinalis’* AOX in human cells and possibly shed some light on its interactions with respiratory chain. Our approach identifies AOX predominantly to localize to mitochondria (Fig. 5a, Supplementary Fig. 6). More specifically, >90% of AOX interactors (Fig. 5b) belong to mitochondria according to CellWhere database and 48% among them have the GO term annotation of mitochondria inner membrane. Furthermore, 38 of the interactors are components of the mitochondrial respiratory complexes I-V. From these AOX prefers interactions with complex II (2/4 components detected), complex I (19/44) and complex V (9/19) (Fig. 5c), which is also visible from the quantitative interactor abundance (Fig. 5d). Therefore, our MS-microscopy findings confirm the functionally suggested location of transgenic AOX in human cells.

**Figure 5:**
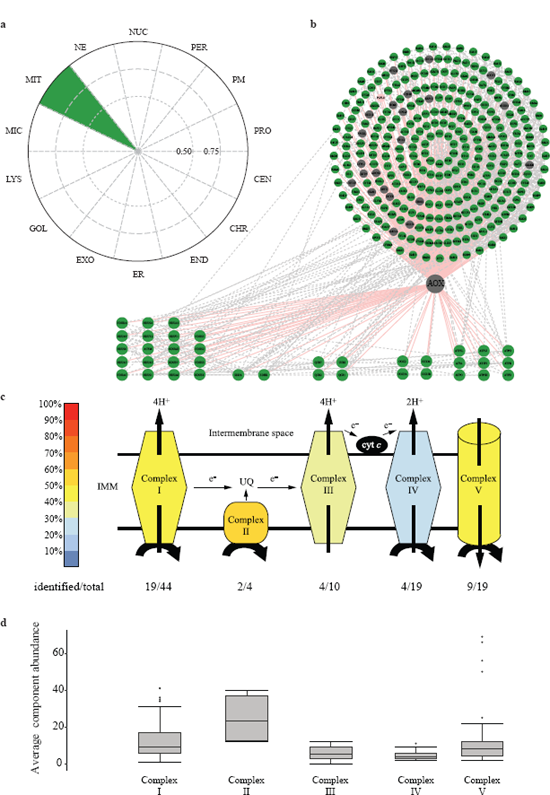
Transgenic alternative oxidase (AOX) localized to the inner mitochondrial membrane facing to the mitochondrial matrix and locating proximal to Complex II. (a) The AOX from *Ciona Intestinalis*, was introduced to human cells and the MS-microscopy assigns the AOX to localize to mitochondria. (b) The BioID approach identifies 333 interactions of which 93.1% (310) were mitochondrial (green), 0.3% (1) was peroxisomal (pink) and 6.6% (22) were unassigned (grey), based on CellWhere database. (c) Total of 38 of the interactors were components of the mitochondrial respiratory chain complexes (Complex IV, key: color gradient indicates the percentage of proteins of the each individual complexes identified). (d) The average component abundance shows that the AOX associates most with the Complex II, which is in agreement with the AOX suggested functional role.

### Defining interaction distances within a protein complex

Others and we have shown that AP-MS offers accurate quantification of complex composition allowing calculations on complex stoichiometry^9,10,40^. With BirA* the labeling radius is limited (circa 10 nm), and it has been used to obtain rough maps of spatial distribution of proteins within structures by reciprocally analyzing BirA*-tagged proteins throughout the structure^15^. As the *in vivo* biotinylation is enzymatic reaction deriving relational interaction abundances of participants cannot be done. However, it can be reasoned that the more proximal proteins will be more efficiently biotinylated and purified in larger abundances than proteins further away^15,41^. On the AP-MS side this would correlate with the likelihood of more abundant interactors being more direct than low abundant, in which the interaction could be mediated by other proteins and the interaction with the bait would be secondary or tertiary etc. Therefore, by blotting both the BioID and AP-MS data, in theory, we could obtain relative distance of the MAC-tagged bait protein to its interacting protein in a complex. For testing this hypothesis, we selected CDK7 and CDK8 for which we have previously identified successfully quantitative complex compositions^25^.

We applied our dual-approach, and with both AP-MS and BioID we could detect the CDK7 interactions with TFIIH core components (Fig. 6a)^42-44^. The size of the TFIIH is estimated to be ~10 nm^42^, which is still within the BirA* biotinylation range and should allow measurement of the CDK7 interaction distances for all of the complex components. Before associating with TFIIH, CDK7 associates and forms cyclin-dependent kinase (CDK)-activating kinase (CAK) complex with two regulatory subunits; cyclin H (CCNH) required for CDK7 activity and with RING finger protein CDK-activating kinase assembly factor MAT1 (MAT) which modulates the substrate specificity of the complex^45^. In agreement, both CCNH and MAT1 are detected as closest to CDK7, followed by ERCC2, ERCC3 and TFIIH1. The ERCC2, TFIIH basal transcription factor complex helicase XPD subunit is the bridge linking the CAK module with TFIIH ring-like core and has been shown to directly interact with TFIIH basal transcription factor complex helicase XPB subunit (ERCC3)^46^ and TFIIH1^47^. CDK7 also has been reported to directly interact with TFIIH 1 (Fig. 6d)^48,49^. Our results are in line with both hypothesis, the evidence suggesting that TFIIH1 and ERCC3 have short inter-distances as well as that they both are close to the CAK module (Fig. 6d, f). The core ring structure of TFIIH composed of TFIIH2-4, CDK7 are located adjacent, having highly similar distance to CDK7 (Fig. 6a, f). Similarly, ERCC5 and TFIIH5 are in longer distances from CDK7, suggesting that they are located on the opposite side of the complex from CAK (Fig. 6a, d, f).

**Figure 6:**
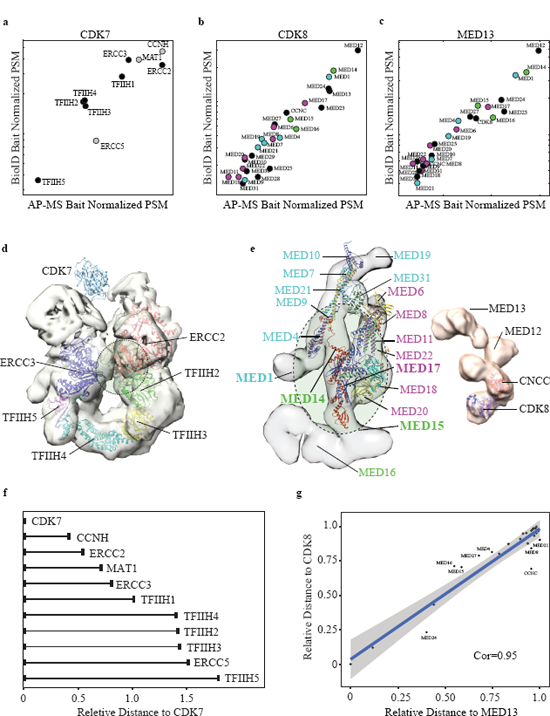
Integration of MAC-tag data allows characterizing of interaction distances within a protein complex. (a-c) Distance based topology of protein complexes. The APMS and BioID data was blotted based on the bait normalized prey abundances and the correlated data was used to derive interaction distances for CDK7 and the TFIIH complex, as well as for CDK8 and MED13 with the Mediator complex. The CDK7 formed CAK-complex components are shown in grey and the Mediator complex components assigned to the Head (magenta), the Middle (cyan) and the Tail (green) are color-coded. (d-e) The derived interaction distances for CDK7, CDK8 and the MED13 are fitted into EM derived complex structures and suggested fitted interaction surface is shown in green dashed line ellipses. The color-coding in e corresponds with the b, c). (f) Relative distances for bait protein and the other complex components can be calculated. (g) The calculated relative distances derived from the integrated APMS and BioID data results to extremely high correlation (c=0.95) for CDK8 and MED13, two neighboring units in the Mediator kinase module.

The transcriptional co-activator Mediator complex has more than 30 subunits and is ~30 nm in size^50^, and therefore on the upper detection limits. The Mediator complex is composed of 4 modules, the head, the middle, the tail and the kinase module. The evolutionarily conserved and dissociable kinase module is formed by CDK8 together with cyclin C (CCNC), mediator of RNA polymerase II transcription subunit mediator complex subunit (MED) 12 and 13, (Fig. 6e)^51-53^. To test and validate the reproducibility of our approach, in addition to CDK8, we additionally choose MED13 for analysis. Additionally this would allow more accurate prediction of the kinase-module docking surface to the Mediator core complex. To our surprise the overall correlation of CDK8 and MED13 distances from the Mediator core is extremely high (c=0.95) (Fig. 6b, c, g), confirming that these two proteins are highly proximal. Based on the analysis the closest Mediator subunits for both CDK8 and MED13 are MED12, MED14, MED1, MED24, MED23, MED17, MED15, MED27, MED16 and MED6/4^53^. This suggests that the kinase module is docking horizontal to the MED14 ranging from RM1 and RM2, the two repeats of a structural domain on MED14.

Both the CDK7, CDK8 and MED13 studies benchmark another utility of our MAC-tag system and shows that by integration of AP-MS and BioID it is possible to derive information on complex structure, interaction distances and possible distance constraints.

## Discussion

In this study, we developed and optimized an integrated workflow based around MAC-tag, for characterization of the molecular context of any protein of interest from human cells. This workflow features state-of-the art affinity purification using Strep-tag to identify and quantify protein-protein interaction and protein complex stoichiometry; identification of transient or close-proximity interactions with BioID; visualization of the bait protein and the proximal interactors with immunofluorescence microscopy; and defining the molecular context with MS-microscopy utilizing the reference dataset obtained by identifying proximal interactors for *bona fide* subcellular localization markers. Additionally our integrated workflow reduces the generation of the required cell lines for APMS and BioID to half.

In addition to analyzing the physical and functional interactions formed by 18 cellular localization markers, we used our integrated workflow to map interactions for four kinases (AURKB, CDK7, CDK8 and GSK3B), as well as for two additional localization markers (BET1 and RAB5A). In addition to identifying 539 interactions for these six proteins, we could validate the accuracy of the MS-microscopy method for identifying correct cellular localization for these proteins. Furthermore, we could show with an exogenous protein, AOX that our MS-microscopy correctly identifies AOX to localize to mitochondria. Additional analysis using our integrated workflow shows that AOX localizes to inner mitochondrial membrane and is in close-proximity with Complex II. Our findings validate, for the first time, the functionally suggested vicinity of AOX with Complex II.

Identifying the complex components in a stoichiometry fashion has been shown to be possible with affinity purification mass spectrometry^9,10^. However, obtaining any further spatial information of the complex formation has only been possibly in combination with XL-MS^18^. We could now show with the TFIIH and Mediator complex as model complexes, that by utilizing both the AP-MS and BioID approaches we can obtain relative interaction distances for proteins in a complex. Based on the interaction distances it is possible to obtain an estimate for the interaction surfaces for proteins or structures, such as with the kinase submodule of the Mediator complex.

In summary, our study showed that the integrated workflow and the reference molecular context proteome map generated here, allows an easy way to probe the molecular localization of protein of interest, and additionally an online resource of our BioID based MS-microscopy approach is available at http. The “molecular image” obtained from the MS-microscopy analysis considers the weights of interactors and provides more dynamic localization information at the molecular level. The developed MAC-tag and the integrated approach should empower, not only the interaction proteomics community, but also cell biologist, with an experimentally proven integrated workflow for mapping in detail the physical and functional interactions and the molecular context of any protein in human cells.

## Methods

### Generation of MAC-tag Gateway^®^ destination vectors

To generate Gateway compatible destination vectors, plasmids *containing* the tags (C terminal: StrepIII/HA/BirA*, N terminal: BirA*/HA/StrpIII) were synthesized by GeneArt^®^, Life Technologies. These were digested with restriction enzymes and inserted into N terminal: pcDNA5/FRT/TO/ StrepIII/HA/GW^19^ or C terminal: pcDNA5/FRT/TO/ StrepIII/HA/GW^2^ in which entire StrepIII/HA tag was removed. All the Gateway compatible entry clones, which contain subcellular marker gene of interested, were from Human ORFeome collection.

### Immunofluorescence

HeLa cells (American Type Culture Collection) were transfected with vectors containing MAC-tagged gene of interest and cultured either with or without supplemental biotin. Bait proteins were detected with anti-HA antibody, followed by Alexa Fluor488-conjugated secondary antibody. Biotinylated proteins were detected with Alexa-Fluor 594 streptavidin. DAPI staining was used to determine the nuclei. Wide field fluoresce microscope (Leica DM6000, Leica) with HCXPL APO 63x/1.40-0.60 oil objective was used to image the samples. The image files were processed with LAS X (Leica), and ImageJ softwares.

### Cell culture

For generation of the stable cell lines inducibly expressing the MAC-tagged versions of the baits, Flp-In™ 293 T-REx cell lines (cultured in DMEM (4.5 g/L glucose, 2 mM L-glutamine) supplemented with 10% FBS, 50 mg/mL penicillin, 50 mg/mL streptomycin) were co-transfected with the expression vector and the pOG44 vector (Invitrogen) using the Fugene6 transfection reagent (Roche Applied Science). Two days after transfection, cells were selected in 50 mg/mL streptomycin and hygromycin (100 μg/mL) for 2 weeks, and then the positive clones were pooled and amplified. Stable cells expressing MAC-tag fused to green fluorescent protein (GFP) were used as negative controls and processed in parallel to the bait proteins.

Each stable cell line was expanded to 80% confluence in 20 × 150 mm cell culture plates. Ten plates were used for AP-MS approach, in which 1 μg/ml tetracycline was added for 24 h induction, and ten plates for BioID approach, in which in addition to tetracycline, 50 μM biotin was added for 24 h before harvesting. Cells from 5 × 150 mm fully confluent dishes (~ 5 × 10^7^ cells) were pelleted as one biological sample. Thus, each bait protein has two biological replicates in two different approaches. Samples were snap frozen and stored at – 80 °C.

### Affinity purification of the interacting proteins

For AP-MS approach, the sample was lysed in 3 ml of lysis buffer 1 (0.5% IGEPAL, 50 mM Hepes, pH 8.0, 150 mM NaCl, 50 mM NaF, 1.5 mM NaVO_3_, 5 mM EDTA, supplemented with 0.5 mM PMSF and protease inhibitors; Sigma).

For BioID approach, Cell pellet was thawed in 3 mL ice cold lysis buffer 2 (0.5% IGEPAL, 50 mM Hepes, pH 8.0, 150 mM NaCl, 50 mM NaF, 1.5 mM NaVO_3_, 5 mM EDTA, 0.1% SDS, supplemented with 0.5 mM PMSF and protease inhibitors; Sigma). Lysates were sonicated, treated with benzonase.

Cleared lysate was obtained by centrifugation and loaded consecutively on spin columns (Bio-Rad) containing lysis buffer 1 prewashed 200 μl Strep-Tactin beads (IBA, GmbH). The beads were then washed 3 × 1 ml with lysis buffer1 and 4 × 1 mL with wash buffer (50 mM Tris-HCl, pH 8.0, 150 mM NaCl, 50 mM NaF, 5 mM EDTA). Following the final wash, beads were then resuspended in 2 × 300 μL elution buffer (50 mM Tris-HCl, pH 8.0, 150 mM NaCl, 50 mM NaF, 5 mM EDTA, 0.5mM Biotin) for 5 mins and eluates collected into an Eppendorf tubes, followed by a reduction of the cysteine bonds with 5 mM Tris(2-carboxyethyl) phosphine (TCEP) for 30 mins at 37 °C and alkylation with 10 mM iodoacetamide. The proteins were then digested to peptides with sequencing grade modified trypsin (Promega V5113) at 37 °C overnight. After quenching with 10% TFA, the samples were desalted by C18 reversed-phase spin columns according to the manufacturer’s instructions (Harvard Apparatus). The eluted peptide sample was dried in vacuum centrifuge and reconstituted to a final volume of 30 μL in 0.1% TFA and 1% CH_3_CN.

### Liquid chromatography-mass spectrometry (LC-MS)

Analysis was performed on a Q-Exactive mass spectrometer using Xcalibur version 3.0.63 coupled with an EASY-nLC 1000 system via an electrospray ionization sprayer (Thermo Fisher Scientific). In detail, peptides were eluted and separated with a C18 precolumn (Acclaim PepMap 100, 75μm × 2cm, 3 μm, 100 Á, Thermo Scientific) and analytical column (Acclaim PepMap RSLC, 50 μm × 15 cm, 2 μm, 100 Å, Thermo Scientific), using a 60 minute buffer gradient ranging from 5 to 35% buffer B, followed by a 5 min gradient from 35 to 80% buffer B and 10 min gradient from 80 to 100% buffer B at a flow rate of 300 nl/min (buffer A: 0.1% formic acid in 98% HPLC grade water and 2% acetonitrile; buffer B: 0.1% formic acid in 98% acetonitrile and 2% water). For direct LC-MS analysis, 4 μl peptide samples were automatically loaded from an enclosed cooled autosampler. Data-dependent FTMS acquisition was in positive ion mode for 80 min. A full scan (200-2000 m/z) was performed with a resolution of 70,000 followed by top10 CID-MS^2^ ion trap scans with resolution of 17,500. Dynamic exclusion was set for 30 seconds. Acquired MS^2^ spectral data files (Thermo. RAW) were searched with Proteome Discoverer 1.4 (Thermo Scientific) using SEQUEST search engine of the selected human component of UniProtKB/SwissProt database (http://www.uniprot.org/, version 2015-09). The following parameters were applied: Trypsin was selected as the enzyme and a maximum of 2 missed cleavages were permitted, precursor mass tolerance at ±15 ppm and fragment mass tolerance at 0.05 Da. Carbamidomethylation of cysteine, was defined as static modifications. Oxidation of methionine and biotinylation of lysine and N-termini were set as variable modifications.

### Identification ofthe HCIs

Significance Analysis of INTeractome (SAINT) express version 3.6.0^20^ and Contaminant Repository for Affinity Purification (CRAPome, http://www.crapome.org/)^21^ were used as statistical tools for identification of specific high-confidence interactions from our AP-MS data. 16 GFP control runs (8 N-terminal MAC-GFP and 8 C-terminal MAC-GFP) were used as control counts for each hit and the final results only considering proteins with SAINT score ≥0.73. This corresponds to an estimated protein-level Bayesian FDR of <0.05. Furthermore, we used the CRAPome database with a cut-off frequency of ≥20% (≥82) except the average spectral count fold change ≥ 3 was set for assigning high confidence interactions (HCIs). Clustering analysis Prey protein frequency count matrix was generated using DAVID gene functional classification tool to provide the gene ontology (GO) terms (domains, biological process and molecular function). The p-values associated with each annotation terms has p< 0.01.

Hierarchical cluster was performed by centered correlation (both baits and interactors; average linkage) using Cluster 3.0 and the clusters were visualized with Tree View 1.1.6 and the matrix2png web server (http://www.chibi.ubc.ca/matrix2png/).

### Networks and maps

Protein interaction networks are constructed from SAINT data that were imported into Cytoscape 3.2.1^22^. The Known prey-prey interaction data were obtained from PINA2 database (http://omics.bjcancer.org/pina/)

### MS-microscopy database construction

The high-confidence interacting proteins (HCIPs) obtained from previous filtering steps were sorted according to the corresponding bait protein localization information to build the reference database, containing the following localization information: peroxisome, microtubule, endosome (combined: early, late and recycling endosome), proteasome, nuclear envelope, Golgi (combined: Trans-and Cis-Golgi), lysosome, nucleolus, plasma membrane, endoplasmic reticulum, mitochondria, centrosome, chromatin, exosome.

### Score calculation

Final localization scores for all localization groups for given bait of interest were calculated by dividing the sum of peptide-spectrum match (PSM) values of the interactors that match between bait of interest and a localization group by the sum of PSM values of all bait interactors that match any localization group. This is then multiplied by the sum of PSM values of interactors of the localization group that match the bait of interest interactors divided by the sum of PSM values of all interactors of the localization group. The score reflects subcellular localization by numerically describing the similarity of the subcellular environment and the time spent there between the bait of interest and each localization group.

## Results visualization

The MS-microscopy analyses are presented as polar plots with in-house python script, where the circle has been equally divided into 14 sectors, each sector representing one specific subcellular location. Different colored sector areas indicates the possible location score of query bait, with scores between 0 and 0.5 marked in red, between 0.5 and 0.75 in yellow, and 0.75 and 1 in green.

### Online interface

We have developed web application (R-shiny; http://www.biocenter.helsinki.fi/bi/protein/msmic) for observation of protein localization by MS-microscopy. A user can upload an input file (after SAINT and CRAPome filtering) and visualize the bait protein dynamic localization. The image as well as a parsed data can also be downloaded.

### Determining relative intra molecular distances of the protein complexes

The PSMs of preys were normalized with interacting bait PSM abundance respectively. The scatter plots were based on the normalized PSMs from both BioID and AP-MS approach. Then the Euclidean distances were calculated between bait to preys according to the scatter plot. Subsequently, the correlation was calculated.

### Data deposition

Mass spectrometry data are available at the Pep-tideAtlas (http://www.peptideatlas.org/) raw data repository (PASS01076). Interactome data will be available at the IntAct database.

## Author Contributions

M.V. and X.L conceived the study and designed experiments. M.V., X.L., K.S. and F.T. performed experiments and data analysis. M.V., X.L., K.S. and F.T. participated in manuscript preparation. M.V. and X.L. wrote the manuscript.

## Acknowledgments

We thank Sini Miettinen for technical assistance and Drs. Tiina Öhman and Salla Keskitalo for critical reading and comments on the manuscript. This study was supported by grants from the Academy of Finland (nos. 288475 and 294173), Sigrid Jusélius Foundation, University of Helsinki Three-year Research Grant, Biocentrum Helsinki, Biocentrum Finland, HiLIFE and Instrumentarium Research Foundation.

---

**Supplementary Figure 1:**
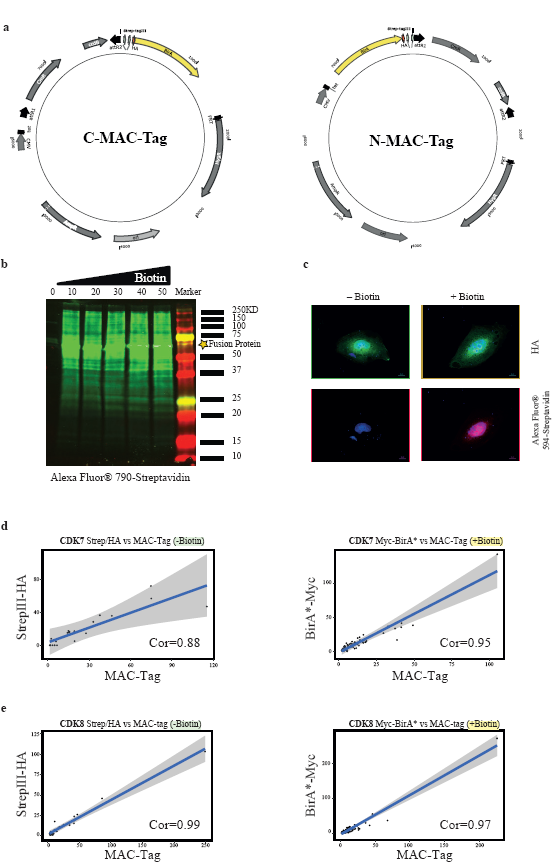
Features of the MAC-tag. (a) Plasmid maps are graphical representation of the MAC-tag vectors. (b) The western blot shows biotinylated proteins of MAC-tagged GFP cell lysate with biotin concentration gradient (0-50 μM) in culture medium by Alexa Fluor^®^594-conjugated streptavidin. (c) Immunofluorescence analysis shows no detectable biotinylation in untreated sample and significant activation of the biotinylation with 10 μM biotin and peaking at 50 μM concentration (anti-HA: green; Alexa Fluor^®^ 594-conjugated streptavidin: red; DAPI: blue. Scale bar: 10 μm). (d-e) There is strong correlation between MAC-tag AP-MS (no biotin) & StrepIII-HA and MAC-tag BioID (with biotin) & BirA*-Myc. A scatter plot shows 95% confidence interval of correlation between MAC-tag data and the data from StrepIII-HA & BirA*-Myc alone.

**Supplementary Figure 2:**
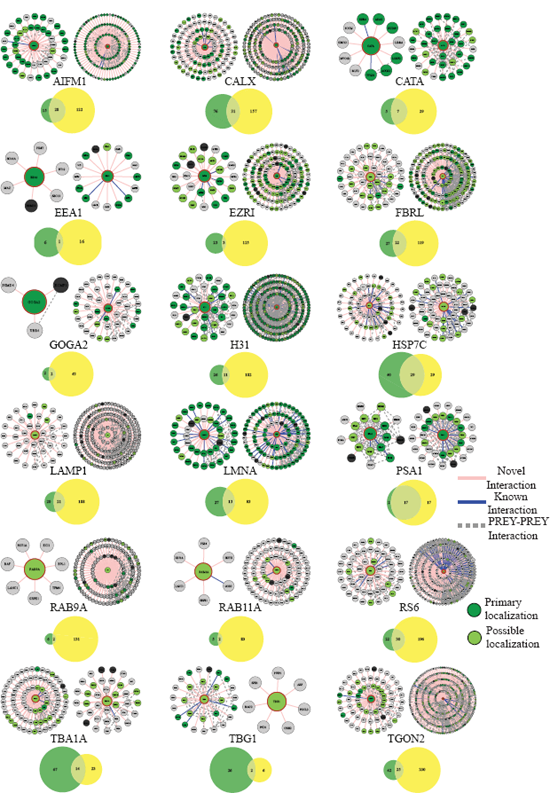
Overview of high-confidence interactions of 18 bona fide cellular localization markers. The individual distribution of interactions detected with of AP-MS and BioID approaches with MAC-tagged 18 *bona fide* cellular localization markers. The novel interactions are presenting in pink lines and the blue line represents the known interaction. Prey-prey interactions represent via dash lines. The nodes are color-coded based on the localization rank obtained from the CellWhere database (key: dark green = primary cellular localization for the corresponding protein, light green = possible localization, grey = different or localization assigned for the protein). Venn’s diagram compares the number of interactions between AP-MS and BioID methods, overlap showing the number of common interactions.

**Supplementary Figure 3:**
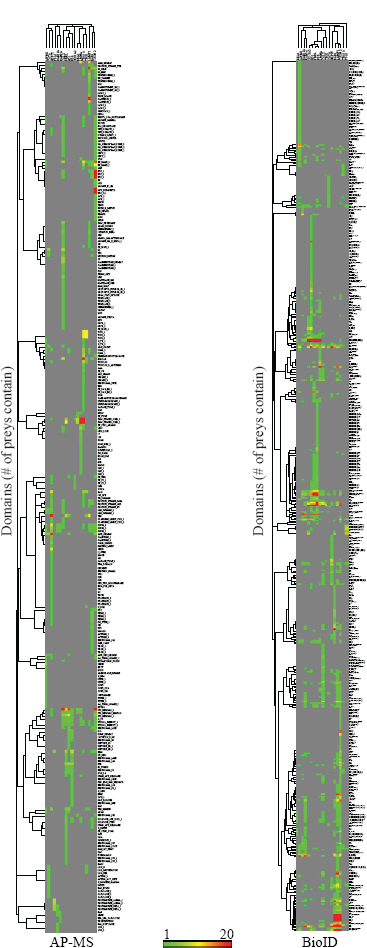
Hierarchical clustering the domain containing of the HCIPs of 18 localization markers. The DAVID bioinformatics recourses 6.8 database (https://david.ncifcrf.gov/) were used for interaction domain containing analysis to generate the value count matrix for clustering (P < 0.01). The color intensities indicate the sum of domain containing count.

**Supplementary Figure 4:**
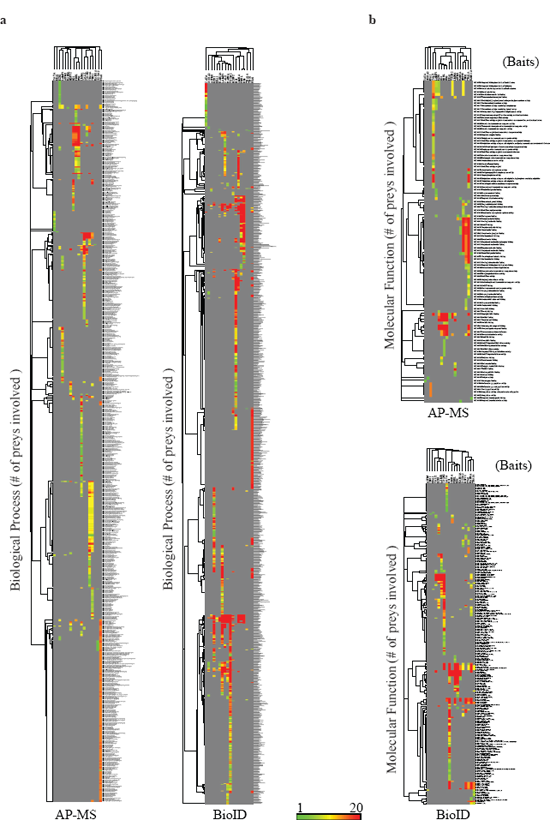
Gene Ontology analysis of interaction contexts of 18 localization markers. The DAVID bioinformatics recourses 6.8 database (https://david.ncifcrf.gov/) are used for GO term analysis to generate the value count matrix for clustering with a P-value cut-off of P < 0.01. The hierarchically clustered heatmap including: biology process (a) and molecular function (b). The color intensities indicate the sum of preys of corresponding GO term. Hierarchal clustering was used to generate the cluster dendrogram.

**Supplementary Figure 5:**
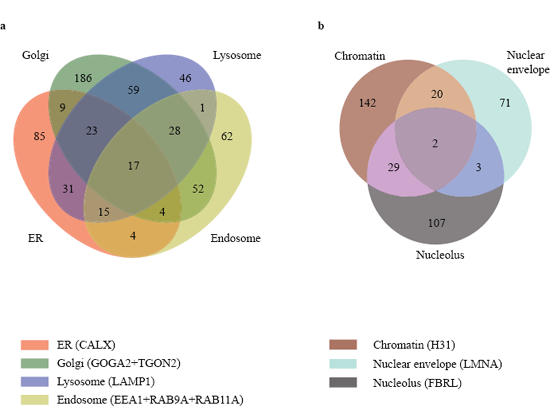
Inter-relation of different subcellular organelles. Venn’s diagram compares the number of identified HCIPs from different subcellular compartments to the biological relationship between different organelles. (a) BioID identified proteins that are trafficking among Golgi, lysosome, endosome and ER are shown. (b) BioID identified proteins trafficking among nucleolus, NE and chromatin are shown.

**Supplementary Figure 6:**
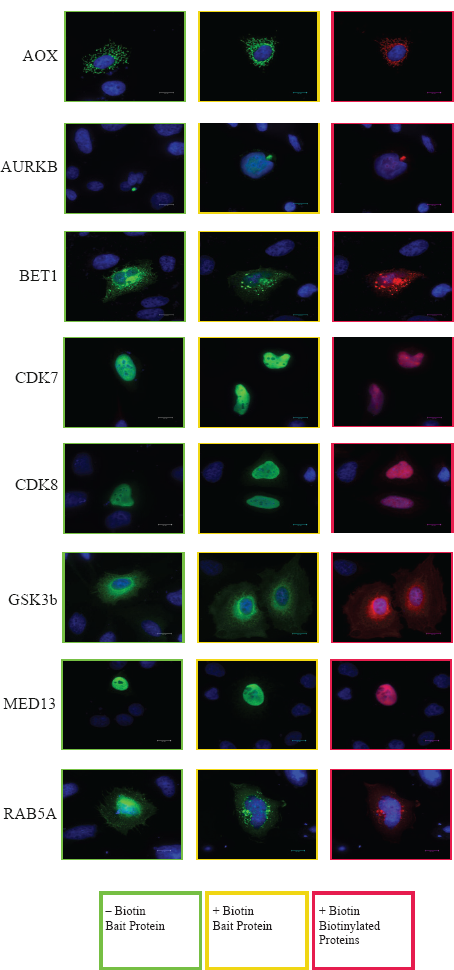
Validation of protein localization with immunofluorescence. Fluorescence microscopy is applied to observe the MAC-tagged proteins that have been monitored by MS-microscopy (Fig. 4). These tagged bait proteins are visualized with anti-HA immunostaining (green), DAPI (blue) and the in vivo biotinylated interactors are staining with Alexa Fluor^®^594-conjugated streptavidin (red), Scale bar: 10 μm.

